## Supplementary Information for "Social and endogenous infant vocalizations"

Appendix A. Focus of prior literature in infant vocalizations…2

Appendix B. Opinion survey on the function of infant vocalizations…4

Appendix C. Expected and actual circumstance durations…8

Appendix D. The origin of vocal flexibility in humans and the fitness signaling hypothesis…10

References. Appendix references…17

### Appendix A

#### Focus of prior literature in infant vocalizations

It has been our impression that most literature in infant and child speech and language development has tended to gather data in interactive circumstances. The work has tended to place primary emphasis on the vocalizations of babies in terms of their vocal interactivity and on contingencies between adult vocalizations and infant responses as well as on adult elicitations and responsivity to infant sounds. These tendencies in the literature, we have surmised, have yielded relatively little attention to endogenously produced infant protophones. Assuming our impression is correct, the literature's tendency is surprising since our data in the present paper suggest most infant protophones are not directed to other persons. But is our impression of the literature consistent with the facts?

In response to a reviewer suggestion we conducted a Pubmed search. We focused on abstracts only. The term "infant vocalization" returned many abstracts that showed no emphasis on social vs. endogenous human infant vocalization, and so were irrelevant to our impression of a primarily social emphasis in the literature. Some abstracts that were returned in the search, for example, merely reported acoustic data on infant sounds, with no mention of either independent/endogenous or interactive/social production. Many were about cry only, not protophones. A great many were not about human vocalization at all (birds were a particular focus). We ignored all such articles as well as articles from or in collaboration with our lab (i.e., with Oller as at least a co-author).

We examined the abstracts for the first 160 articles returned by the search (dates of the 160 articles ranged from 2014 to 2020); only 18 of them could be judged with moderate certainty based on the abstracts regarding having an emphasis on social use of human speech-like vocalizations as opposed to endogenous or independent use. None of these 18 appeared to have attempted to address the question of the current paper (actually counting and focusing on a comparison of rates of social and endogenous use of infant vocalization). Also none seemed to have placed primary emphasis on independent, endogenous production. Fifteen revealed a focus on the social-interactive use of infant sounds, while 3 described use of infant sounds when infants were not interacting, though consideration of interaction was not excluded in these cases.

So all in all, the review seemed to suggest our impression of lack of emphasis on endogenous protophone production in the literature is essentially accurate.

### Appendix B

### Opinion survey on the function of infant vocalizations

As background information for the primary goal of this research, we sought survey data where both parents and non-parents were asked to provide estimates of how often they thought infants vocalize with social directivity and without social directivity based solely on a reflection of their own experiences around infants. We hypothesized that survey participants would provide evidence supporting our general impression of the literature on vocal development, an impression suggesting that socially-directed vocalization is emphasized more often than endogenous vocalization.

#### Materials and Methods

We collected survey data using Amazon Mechanical Turk ("mTurk") to provide a perspective on the observational data in the main text of the article and an empirical evaluation of the suspicion that not only many researchers in child development, but also the general public, have the impression that infants predominantly vocalize socially. mTurk is increasingly used as an online recruitment tool for participation in experimental studies and academic surveys as a quick method to obtain many responses from the general public. mTurk has been shown to be slightly more representative of the US population than of other countries and is considered to be as reliable as traditional survey methods [1–3]. mTurk qualifications used for this study included: 1) having a HIT Approval Rate greater than 95% and 2) at least 50 Approved HITs. These ratings ensured that all participants were experienced and had been deemed acceptable participants in prior mTurk studies. Such qualifying indicators are regularly used by mTurk researchers to safeguard against inaccurate and inattentive workers.

#### Survey instructions

Following consent, participants were presented the following written instructions for the survey:

*This is a study evaluating your perception of how often babies make different kinds of sounds and why they make them. You will be asked to consider sounds produced by babies at three different ages: Infants who are 3-months, 6-months, and 10-months old. Across any given day, consider all the sounds (or "vocalizations") babies make. Your task is to estimate the percent of these sounds that serve a particular function (social or endogenous). In answering the questions, consider your previous experiences (if any) around babies and give an intuitive guess for each question. When thinking about your responses, only consider babies who are typically developing, not those who may have special conditions causing atypical development. You are not expected to be an expert on this, and there are no wrong answers. You will be asked to give an intuitive response. Your responses will be required to sum to 100 (e.g., 100%).*

Participants estimated a percentage of social and endogenous infant vocalization functions at three ages (3-month-olds, 6-month-olds, and 10-month-olds) for a total of six judgments. Means and standard deviations of these responses were calculated. The data in the main text based on laboratory-coded observations of real infants in audio-video recordings were collapsed to yield the same categories (social vs. endogenous) for each infant utterance. The coding in the laboratory as reported in the main text was based on a more extensive list of "illocutionary categories" including No Force (no discernible social force, but also not obviously playful), Vocal Play (not directed to a person or object, but apparently playful), Object Directed, Complaint (this is would be a protophone suggesting distress by not directed to any listener), Directed Complaint (distressful protophone directed to another person), Exultation (celebratory protophone not directed to another person), Directed Exultation (celebratory protophone directed to another person), Continue Interaction (protophone that responds to an adult vocalization or continues in vocalizing during an interaction), Call Initiate (protophone produced by the infant in an apparent attempt to initiate interaction), and Imitation (a protophone that appears to be not only response to an adult vocalization but to include any perceived imitative feature of the adult vocalization). No Force, Vocal Play, Object Directed, Complaint, and Exultation were collapsed for the analysis in the main text to the category "Endogenous", while all the other categories were collapsed to "Social" [4].

#### Survey participants

300 participants completed the online survey, and 239 participants' data were used in final analysis based on adequate responses to three attention checks distributed throughout the survey. The attention checks ensured that the responders were not robots and that the responders

were sufficiently knowledgeable in English to have understood the questions clearly. The attention checks were questions presented to all participants:

1) *Provide a word that means the opposite of happy.*
2) *Select the option that includes 5 times a week:*
    a. *one time a week,*
    b. *two to three times a week,*
    c. *four to five times a week,*
    d. *six to seven times a week.*
3) *Type in the number sixty.*

For the first attention check, a variety of English words meaning " not happy" were accepted. For the second attention check, options C and D were accepted. For the third attention check, only the number "60" was accepted. Failure on two of the attention checks resulted in rejection of the participant. In general, such questions capture robots, which fail usually to answer the questions in a meaningful way. For language background, the participants were asked to list the languages they speak in the order of most to least fluent. Only individuals indicating that English was at least second on their list were included. An additional measure to try to limit the group to English speakers was the inclusion of a worker qualification in the mTurk survey settings that required the computer system location to be in one of the following countries where English is the primary spoken language: AU, CA, NZ, GB, or US. In other words, the worker had to reside in or be taking the survey on a computer registered in one of these countries. Detailed demographics of the mTurk survey participants are presented in Table B1.

#### Results

Fig B1 shows the survey participants' distribution of responses on relative percentages of protophones across the three ages. On average across the three ages, the respondents thought approximately 43% of infant protophones were endogenous. In addition, they thought infants produce fewer endogenous vocalizations at the end of the first year (36%) than at the beginning (50%). Thus, the respondents believed more than half of infant protophones are socially directed and many more than half by 10 months.

**Table B1. mTurk survey participant demographics.** Participant demographics for opinion study.

| **Age** | | **Gender** | | **Education** | | **Number of children** | | **Frequency around children** | |
|---|---|---|---|---|---|---|---|---|---|
| **18-21** | 3 | **Male** | 139 | **Less than HS** | 2 | **None** | 124 | **Never** | 29 |
| **21-34** | 126 | **Female** | 97 | **HS/GED** | 29 | **1** | 41 | **Rarely** | 83 |
| **35-44** | 50 | **Other** | 3 | **Some college** | 48 | **2** | 41 | **Sometimes** | 62 |
| **45-54** | 34 | | | **Associate's** | 33 | **3** | 21 | **Frequently** | 46 |
| **55-64** | 24 | | | **Bachelor's** | 111 | **4+** | 12 | **All the time** | 19 |
| **65+** | 2 | | | **Master's** | 9 | | | | |
| | | | | **Doctorate (PhD)** | 2 | | | | |
| | | | | **Professional Degree (JD, MD)** | 5 | | | | |

**Fig B1. mTurk opinion study on social directivity of infant protophones across 3 ages.** Opinions of the survey participants on how often infants use protophones socially and endogenously. Participants believed infants decrease the percentage of endogenous protophones between 3-10 months, from 50% at the youngest age to 36% by the oldest age.

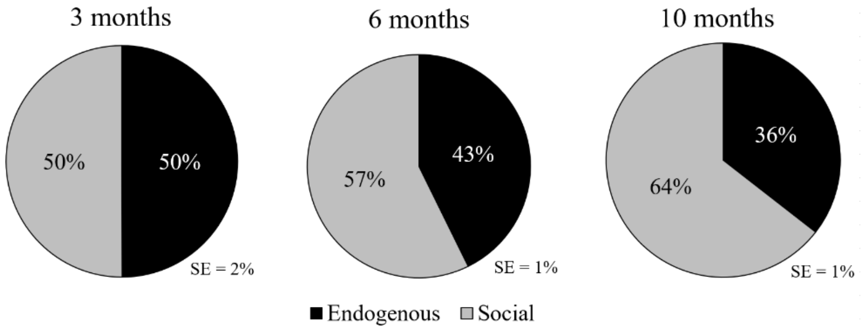


Parents and non-parents reported similar percentages of social and endogenous vocalizations. Overall, parents reported infants used protophones socially 58% of the time, whereas non-parents reported 57%. Males and females also estimated very similar percentages of social protophones (58 and 57% respectively). Persons who self-identified as being around kids "all the time" estimated that infants produce 58% social protophones, while those who self-identified as never being around kids estimated 55%. For all these comparisons (parents v. non-parents, males v. females, always around kids v. never around kids), the estimated percentage of social protophones was higher at 6 than 3 months and higher at 10 than 6 months.

### Appendix C

#### Expected and actual circumstance durations

During recording, parents were asked to participate in two recording protocols. For twenty minutes each, parents were asked to engage in face-to-face interaction with their infant (parent seeking to be Engaged with the infant) and to converse with an interviewer while the infant was in the room (parent and infant Independent). Each protocol was initially labeled per the “expected” session protocol. These expected protocol durations are presented in Table C1. The times varied because infant state varied, and sessions were often readjusted for length to keep infants comfortable and often because parents requested the readjustments.

**Table C1. Expected protocol durations.** Duration of expected protocol sessions in Engaged (Engd) and Independent (Ind) protocols for each infant at each age. The minimum duration was 12:08, maximum duration 22:03, with an average duration of 19:10.

| Infant | Gender | Length of recording | | | | | |
|---|---|---|---|---|---|---|---|
| | | Engd | Ind | Engd | Ind | Engd | Ind |
| 1 | F | 00:19:41 | 00:14:13 | 00:18:42 | 00:19:29 | 00:19:58 | 00:19:58 |
| 2 | M | 00:20:03 | 00:20:19 | 00:20:47 | 00:21:05 | 00:21:16 | 00:20:24 |
| 3 | M | 00:22:03 | 00:22:01 | 00:20:20 | 00:20:13 | 00:20:11 | 00:12:52 |
| 4 | F | 00:19:01 | 00:19:39 | 00:16:03 | 00:19:37 | 00:19:51 | 00:19:52 |
| 5 | M | 00:16:11 | 00:19:51 | 00:20:54 | 00:18:11 | 00:20:52 | 00:20:54 |
| 6 | F | 00:19:48 | 00:19:53 | 00:12:08 | 00:14:23 | 00:19:08 | 00:19:54 |
| **Mean age** | | **3 months** | | **6 months** | | **10 months** | |

To encourage naturalistic interaction throughout all protocols, parents were not restricted from engaging with the infant or another person if warranted despite the expected protocol (e.g., to comfort the infant if crying during the *Independent* protocol or to answer a question from a staff member during the *Engaged* protocol). Because the parent would occasionally engage with the infant for notable periods of time during the “expected” *Independent* protocol, or to converse with a staff member during the “expected” Engaged protocol, each recording was re-coded, segmenting it into “actual” *Engaged* and *Independent* circumstances. These periods of time were summed at each age for each infant to create actual protocol durations, shown in Table C2. Four cells in the Table (for Infants 1 and 6 at three and six months in *Independent* circumstance) showed actual protocol durations of less than 5 minutes each, highlighted with an (*).

**Table C2. Actual protocol durations** Duration of actual segments concatenated with *Engaged* (Engd) and *Independent* (Ind) activity for each infant at each age. Overall, there were longer periods of time in the *Engaged* circumstance compared to the *Independent* circumstance. The minimum duration was 00:58, maximum duration 32:52, with an average duration of 19:06.

| Infant | Gender | Length of recording | | | | | |
|---|---|---|---|---|---|---|---|
| | | Engd | Ind | Engd | Ind | Engd | Ind |
| 1 | F | 00:32:38 | 00:01:16* | 00:33:48 | 00:04:23* | 00:20:34 | 00:19:22 |
| 2 | M | 00:27:59 | 00:12:24 | 00:26:59 | 00:14:53 | 00:23:34 | 00:18:08 |
| 3 | M | 00:22:46 | 00:21:19 | 00:23:08 | 00:17:28 | 00:25:35 | 00:07:29 |
| 4 | F | 00:23:26 | 00:15:15 | 00:10:31 | 00:25:08 | 00:24:27 | 00:15:16 |
| 5 | M | 00:22:00 | 00:14:02 | 00:20:54 | 00:18:11 | 00:21:45 | 00:19:55 |
| 6 | F | 00:35:52 | 00:01:37* | 00:25:33 | 00:00:58* | 00:24:02 | 00:15:00 |
| **Mean age** | | **3 months** | | **6 months** | | **10 months** | |

A ratio of expected over actual times for each circumstance and age is presented in Table C3. For most circumstances, larger ratios are seen in *Engaged* circumstances, showing parents were often inclined to engage with their infant in both expected *Engaged* and expected *Independent* circumstances, often running counter to the protocol instructions.

**Table C3. Ratio of expected over actual.** Ratios for actual over expected protocol durations in *Engaged* (Engd) and *Independent* (Ind) circumstances. Larger ratios are seen in the *Engaged* circumstances for all but two infant ages (Infants 4 and 5 at six months).

| Infant | Gender | Length of recording | | | | | |
|---|---|---|---|---|---|---|---|
| | | Engd | Ind | Engd | Ind | Engd | Ind |
| 1 | F | 1.66 | 0.09 | 1.81 | 0.23 | 1.03 | 0.97 |
| 2 | M | 1.40 | 0.61 | 1.30 | 0.71 | 1.11 | 0.89 |
| 3 | M | 1.03 | 0.97 | 1.14 | 0.87 | 1.27 | 0.58 |
| 4 | F | 1.23 | 0.78 | 0.66 | 1.28 | 1.23 | 0.77 |
| 5 | M | 1.36 | 0.71 | 1.00 | 1.00 | 1.04 | 0.95 |
| 6 | F | 1.81 | 0.08 | 2.11 | 0.07 | 1.26 | 0.75 |
| **Mean age** | | **3 months** | | **6 months** | | **10 months** | |

### Appendix D

#### The origin of vocal flexibility in humans and the fitness signaling hypothesis

Oller and various colleagues, including Long and Bowman (and especially Ulrike Griebel), have written elsewhere on the idea that human development provides key information about likely sources of the selection pressures that have driven hominins to differentiate dramatically from our ape cousins in vocal communication [5–8]. We largely share this reasoning with J. L. Locke who formulated a similar proposal independently [9,10]. In this evolutionary developmental biology or "evo-devo" framework [4,11–13] we have formulated a natural logic of development and evolution, where it is proposed that foundational communicative capabilities must develop in order for subsequent capabilities (ultimately required for language) to be possible. Within that reasoning, an essential foundation for language evolution and human linguistic development is a flexible system of expression, where *all* the elements (the vocal modality has proven to be preferred) can be produced with *any* illocutionary intent (any function). One of those possible intents had to have been exploration of vocalization itself, for no social purpose. Another would have been emotional expression, whether of positive or negative emotions. And another, of course, would have been (or would have developed quickly to become) social interaction involving sharing of emotional states and information (going beyond purely manipulative functions such as pleading for help, a kind of function that is common in mammals). Crucially we have posited that human vocal social interaction is itself founded in flexible vocalization. According to the reasoning, one cannot flexibly share states and information vocally, but can only engage in manipulative vocal interactions (threatening, courting, soliciting…, the kinds of vocal interactions seen in mammals in general), unless one has the flexibility to express states that are *not* bound to particular manipulative goals.

The human infant appears to have such a vocal capability from birth [14,15], producing ~3500 protophones daily [15]. But the other apes appear to have no such capability. In 1700 minutes of longitudinal observation of 3 bonobo infants with their mothers we found not a single instance of a "protophone-like" sound produced by a bonobo infant that was interpreted by the coders as "exploratory" or "playful" [16]. All the "protophone-like" sounds produced by the bonobo infants that could be interpreted for function/affect were interpreted as negative/complaint or plea-like vocalizations (the infant seeking help from the mother or simply complaining).

Importantly we also found not a single case of a maternal vocalization directed to one of the bonobo infants. The mothers were very responsive to the infant pleas, but never vocally. It appears chimpanzees are similarly constrained in vocal interaction with infants [17]. The evidence is consistent with the proposed natural logic: In the absence of flexible vocalization on the part of the infant, there is no basis for development (or evolution) of flexible vocal interaction.

So the fundamental question becomes, what selection pressures could have resulted in a flexible vocalization capability before language existed, indeed before vocal social interaction in apes (not obligatorily manipulative in any particular way or limited to a specific single goal) existed? The results of selection pressures had to have been advantageous to the individuals subject to those pressures at the time they first appeared. Thus they could not have been selected as preparation “for language” or language-like communication because language or language-like communication did not yet exist. This is where the fitness signaling idea has traction. Hominin infants, who were more altricial than other ape infants, were more in need of parental care and for a longer period of development than other ape infants. In accord with the proposal, hominin infants were thus under heightened selection pressure to signal their wellness, and vocalization became one of the targeted means of doing that.

Hominin infants were, then, selected to produce protophone-like sounds endogenously and flexibly, especially in circumstances of comfort and lack of immediate need, because in that way, caregivers could recognize and judge the wellness of the infants. The advantage to the caregivers was greater efficiency in their investments in offspring, yielding presumably more numerous progeny in subsequent generations. Hominin infants are thus seen as having been in competition with each other for parental investment and so were selected generation after generation to be increasingly inclined to vocalize in a variety of states including in comfort and with illocutionary flexibility, that is, to produce protophones. The availability of these flexible sounds, recognized by caregivers (who were themselves, according to the proposal, under selection pressure to accurately recognize the well-being of their infants) afforded the opportunity for comfortable vocal interaction among parents and infants (a type of interaction largely absent in other apes, with the exception of certain “close calls” that in some cases occur during grooming), during which parents had further opportunity to observe and even elicit vocal fitness signaling. The flexible vocalizations of the ancient hominin infants provided the raw material of vocalization for parent-infant vocal interactions. The parents of the infants, having been the beneficiaries of the same selection pressure on vocal flexibility from their own infancies, are imagined within the proposal to have developed further vocal flexibility as they matured, along with increasing interest in observation of their infants’ vocal capacities as information about their fitness.

Bonding and attachment of hominin parents and infants seem to have come to be pursued in part during and through these flexible and comfortable vocal interactions. Generation by generation the infant tendency to vocalize freely and the parental tendency to intuitively recognize the import of the protophones grew, according to the proposal, cyclically, ratcheting up the vocal capacities and vocal interactions of hominins across the life span and forming foundations for additional vocal communicative growth.

The fitness signaling function of the protophones did not require that the sounds be intended by the infants as fitness signals—the perlocutionary effect, that is the reaction of the parent in interpretation of the protophones *as* fitness signals needs to be distinguished from the illocutionary intent of the infant in producing the protophones. The infant’s intent had to be variable on different occasions (or the vocal capacity would not have been functionally flexible), and crucially, at least some of the time that intent had to be purely exploratory, the infant expressing interest in the sound production itself, while on other occasions the same sounds had to be producible as expressions of varying emotional states or in seeking to engage or maintain interactive engagement with a caregiver.

The reason the parent’s reaction needs to be distinguished from the infant’s intents, is in part that regardless of the infant’s intent with protophone production, the parent’s interpretation would affect the parent’s decisions about investment. The parent’s interpretation of the infant’s fitness would have resulted from the protophone production (and of course many other signs of fitness of the infant including body movement, skin color, eye contact…) even if the infant had not intended the sounds as fitness signals. Of course there is no reason to doubt the infant could at least on some occasions produce protophones indeed for the purpose of soliciting parental social attention, and in that way may have intentionally been seeking investment. But not always, and that is the key point. The hominin break from the ape background depended, according to our reasoning, on selection for infants who had both the capability and the inclination to produce all the protophone types (on different occasions) in *any* state and with *any* intention. We have striven to emphasize in all our writings about this point that language in all its forms requires this kind of flexibility of expression, as revealed by the fact that every word or sentence of any language can be produced in any circumstance of state or intention. Put another way, humans can utter any word or sentence with any chosen illocutionary force. The fact that human infants from the first month of life appear to be able to do the same with protophones suggests to us that a fundamental break from the ape background occurred when hominin infants were selected to produce protophones in any state and with any purpose. We reason that parental selection of infants based on their interpretation of protophones as fitness signals resulted, generation after generation, in infants inclined to produce such sounds more and more copiously.

Importantly, the proposal does not suggest that the selection pressures on this system of infant endogenous protophone production and parental interest in those sounds and elicitation of them would have abated in modern times. It is an empirical question how and to what extent the pressures apply nowadays (we are already engaged in studies of behavioral responses of parents and other adults listening to infant sounds and are planning physiological studies of adult responses as well).

In the societies we ourselves have studied, parental attention to infants in the first year tends to be intense, although it varies, for example, by socio-economic status. But even in the circumstance where parents are involved relatively little in interaction with their infants, we have never observed a human infant who did not produce massive numbers of protophones. A key unresolved empirical matter concerns the extent to which modern infants produce protophones in societies where, for example, infant mortality is high, and where there is parental resistance to interacting vocally with very young infants. Resistance even to naming infants until they have proven their survivability has been invoked as a possible corollary of such resistance to vocal interaction with infants in at least some societies. We know of no direct studies of protophone rates produced by infants in such societies, although there are empirical reports suggesting much reduced levels of IDS (see [18]).

Our rationale, then, is built on our proposal that more attention in human development research needs to be directed to the endogenously-produced protophones. That the majority of them seem to be produced without social directivity is surprising to us, and we presume it will be surprising to most readers.

Our methods depend on coders' acting as intuitive observers, noticing moment by moment the direction of infant attention, and taking stock of the fact that infants often direct their attention away from interaction even during periods where the parent is eliciting it, and where the infant is intermittently participating actively in it. As noted by Maya Gratier in her review of a previous version of this work, one should not underestimate the potential importance of those occasions (even if they are relatively rare) where the infant is indeed fully engaged and vocalizes in harmony with the caregiver, that is, where the active infant applies its endogenous capacities in directed dyadic communication. Indeed there is reason to suspect that those events of very engaged face-to-face vocal interaction are critical in social and language development. The present paper emphasizes that the infant's endogenous vocalization provides raw material that is required for such active, comfortable vocal interaction.

Furthermore the endogenous vocal tendencies of infants appear to play a significant role in the development of the vocal capacity itself—vocal exploration may serve as a sort of practice in phonatory and articulatory skills that not only provide fitness signals but at the same time lay groundwork for subsequent vocal expression. Note again, that the initial selection pressures that have driven the production of protophones in hominins would have had to involve advantages applying *before* linguistic communication existed. Consequently it seems selection pressure on a practice function could not have operated in isolation but would have been logically dependent upon other selection pressures to establish primitive flexible communication through vocalization in the absence of a language target. The fitness signaling function appears to provide a selection pressure that could have operated in the absence of modern language or even of primitive protolanguage.

We are unaware of other proposals that could explain the initial break with the vocal communication limitations of our ape ancestors (although our own proposal is shared by J. L. Locke). It has been suggested we consider alternative proposals regarding the origin of vocal language such as those of Falk [19] and Dissanayake [20]. There are quite a few psychologists, linguists, biologists, and cognitive scientists that we could add to such a list (e.g., [21–25]) But as far as we know, none of these proposals offers an explanation for the initial break from the ape background with regard to vocal flexibility, the event that we think is a prerequisite to all the other requirements that would have had to evolve for language to have ultimately emerged (infant-directed speech, vocal imitation, learned vocal performatives, primitive syntax, and so on).

A proposal of Robin Dunbar is perhaps the most closely related to our own [26–28]. He has argued that vocalization in ancient hominins may have assumed a role similar to that of grooming as hominin group sizes increased and there was insufficient time in the day to physically groom all the necessary members of the group. “Vocal grooming” (which was posited even earlier by Morris [29]) could service multiple members of the group simultaneously. The grooming function was thought to provide a platform for elaboration of human vocalization in subsequent evolution. That close calls (and lip smacks, see [30]) occur sometimes in primate grooming suggests there may have existed a comfortable social function for some vocalizations in our distant ancestors. In addition, protophones, produced in interactive circumstances, can be thought of as a kind of vocal grooming. But the Dunbar proposal does not incorporate the evo-devo perspective, wherein it is assumed that new vocal capacities would have likely been selected for first in infants, whose subsequent development could have laid the groundwork for the occurrence of even more elaborate vocalizations in grooming adults and in other kinds of interactions among adults.

In our opinion, the grooming hypothesis of Dunbar also requires that the earlier question be answered: How might infant vocalizations with functional flexibility (including the flexibility to have been used in grooming) have been selected for before language existed or before other elaborate forms of vocal interaction existed? We propose that fitness signaling offered the opportunity for selection of infants with greater inclination to vocalize flexibly, and once that greater inclination was in place, an important consequence could have been the development of capabilities yielding social grooming and other functions in infancy and in adulthood.

Our theoretical inclination is based 1) on the proposed natural logic of how a language capability could in principle evolve (vocal flexibility is required for all the features of vocal language and language use) [8], 2) on a common tenet of evo-devo, wherein it is assumed and observed that natural selection tends to target development; if a new structure or capability is to emerge, its genetic foundations must be targeted; minor genetic changes can produce significant changes in structures or capabilities through epigenetic, self-organizational development, and 3) on the fact that protophones have been observed to occur copiously in all human infants long before language and in fact months before many of the presumed prerequisites to language. Thus we propose minor genetic changes in ancient hominins could have resulted in greater flexible vocal activity in hominin infants, and that that greater activity could have had cascading consequences on later capabilities relevant to vocal communication.
